# Model-free analysis in the spectral domain of postmortem mouse brain EPSI reveals inconsistencies with model-based analyses of the free induction decay

**DOI:** 10.1101/2022.02.24.481824

**Authors:** Scott Trinkle, Gregg Wildenberg, Narayanan Kasthuri, Patrick La Rivière, Sean Foxley

## Abstract

**Purpose:** Dysmyelinating disorders lead to abnormalities in myelin structure that produce detectable effects in an echo-planar spectroscopic imaging (EPSI) signal. To estimate the voxel-wise proportion of myelin, data are typically fit to compartmental models in the time domain. This work characterizes limitations in these models by comparing high-resolution water spectra measured in postmortem fixed mouse brains to spectra predicted from time-domain models fit to the same data, specifically by comparing spectra from control and shiverer mice, a model for dysmyelination.

**Methods:** Perfusion-fixed, resected control (*n* = 5) and shiverer (*n* = 4) mouse brains were imaged using 3D EPSI with 100 µm isotropic resolution. The free induction decay (FID) was sampled every 2.74 ms over 192 echoes and Fourier transformed to produce water spectra with 1.9 Hz resolution. FIDs were also fit to two biophysical models and the resulting fits were converted to spectra with a Fourier transform. Spectral asymmetry was computed and compared before and after fitting the data to models.

**Results:** Spectra derived from both models did not show the magnitude of asymmetric broadening observed in the raw data. Correlations between data- and model-derived asymmetries and estimated frequency shifts are weak, leading to a reduction in spectral sensitivity to changes in white-matter structure after fitting the data to models.

**Conclusion:** The results demonstrate spectral inconsistencies between biophysical model predictions and measured data, promoting the further incorporation of spectral analysis methods to develop and benchmark new model-based approaches.

## 1 Introduction

Myelin is a lipid-rich substance produced by oligodendrocytes in the central nervous system that physically surrounds axons in order to improve the transmission of action potentials (1). Its importance to the normal function of the human brain has been demonstrated through the symptoms of demyelination disorders such as multiple sclerosis and additional disorders related to defective myelin structure such as hypomyelination, dysmyelination, and myelinolysis (2).

Conventional clinical MRI approaches to the diagnosis of myelin disorders use combinations of *T*_1_-weighted, *T*_2_-weighted, and FLAIR acquisitions and have been shown to have poor specificity to myelin (3– 5), prompting the development of approaches such as myelin water imaging (6, 7), which uses a multi-echo Carr-Purcell-Meiboom-Gill (CPMG) sequence to measure high-fidelity *T*_2_ decay curves at each voxel. Water protons within the myelin sheath have a very short *T*_2_ value (∼20 ms) (8) relative to intra-axonal and extracellular water protons, allowing for the estimation of the myelin water fraction (MWF) from the *T*_2_ distributions at each voxel.

Recently, the use of a multi-gradient echo (MGE) sequence, also referred to as echo-planar spectroscopic imaging (EPSI), has emerged as an alternative to myelin water imaging (9–16). EPSI measures some portion of the voxel-wise 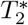 decay curve (the free induction decay, or FID) and results in clinically relevant advantages over the CPMG approach such as faster scan times, a larger volume coverage, and a lower specific absorption rate (17).

The phospholipids that make up the myelin sheath have a magnetic susceptiblity (18, 19) with a diamagnetic property relative to the surrounding water. In the presence of a strong magnetic field, this susceptibility causes a distinct magnetic field perturbation (20) that can be observed as a frequency shift in the spectral domain of the EPSI signal (21). Additionally, voxel-wise changes in FID phase and 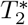 have been shown to depend on the orientation of the white matter fibers relative to the main magnetic field (*B*_0_) (22–26). Evidence from biophysical modeling (18, 27, 28) and experiments (21, 28, 29) suggests that this orientation-dependence is driven by anisotropy in the magnetic susceptibility of myelin (18, 24, 25, 30).

Accordingly, to estimate myelin content and integrity from the EPSI signal, FID curves are typically fit to a biophysical model that assumes the white matter is composed of three distinct, nonexchanging water components: myelin water, intra-axonal water, and extracellular water, each with distinct 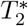 values and potential magnetic susceptibility-dependent frequency shifts (9, 12, 13, 17, 31). While these models are typically fit to FID curves in the time domain, we have shown in previous work (15) that the microstructural features are also observable in a spectral domain representation of the same data after a simple Fourier transform. In the spectral domain, typical models predict that the signal is given by a superposition of three peaks with 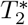-dependent broadening and that the susceptibility-dependent frequency shifts of the underlying components causes the broadening of the overall line-shape to be asymmetric in white matter. Using a model-free approach, we quantified this asymmetric broadening of the water resonance in white matter with a simple asymmetry metric and demonstrated that it is sensitive to the principal orientation of the constituent fibers and *B*_0_.

In an additional study, we performed further spectral asymmetry analysis through a comparison of data from both control and mutant “*shiverer* “ (Mbp^Shi^) postmortem, fixed mouse brains (16). The shiverer model is frequently used to explore MR sensitivity to myelin (32–35) since the mutation causes deletion of most of the myelin basic protein gene (36, 37). This mutation leads to the production of sparse myelin sheaths that are thin or completely absent (38, 39), but with no additional axonal degeneration or inflammation (40). These well-characterized microstructural differences in white matter between the two mouse models have been shown to be a source of difference in the 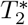-weighted MRI signal (15, 16).

In this work, we extend our analysis of water spectra from control and shiverer mouse brains to explore the performance of two biophysical compartmental models published by Van Gelderen et al. (31) and Nam et al. (12). Specifically, we fit both models to voxel-wise FIDs in the time domain and perform a Fourier transform in time to compare the asymmetry of the water spectrum predicted by the estimated model parameters to the asymmetry observed directly in the data itself. Given the linearity of the Fourier transform, the component-specific frequency shift information computed by the models must be consistent with frequency information resolved in the spectral domain. Accordingly, spectral-domain analysis serves as a direct and convenient method to validate the results of temporal fitting algorithms. To that end, the aim of this work is to apply the models as published and identify discrepancies between the predicted and measured data in the spectral domain that can help improve current models and motivate the development of new models.

## 2 Methods

### 2.1 Sample preparation

Procedures for the collection of the EPSI and diffusion MRI data used for this study have been published in a previous study (16) and are repeated here for completeness. All procedures performed on animals followed protocols approved by the Institutional Animal Care and Use Committee and were in compliance with the Animal Welfare Act and the National Institutes of Health Guide for the Care and Use of Laboratory Animals. Adult mice were deeply anesthetized with 60 mg/kg pentobarbital and sacrificed by intercardial perfusion with a solution (pH 7.4) of 0.1 M sodium cacodylate and heparin (15 units/ml). This was immediately followed by a solution of 2% paraformaldehyde, 2.5% glutaraldehyde, and 0.1 M sodium cacodylate (pH 7.4). Brains were carefully removed from the skulls and post-fixed in the same fixative overnight at 4°C. Brains were soaked in phosphate buffered saline (PBS) prior to imaging for at least 72 hours to remove fixative from the tissue.

### 2.2 MR imaging

Resected control (*n* = 5) and shiverer (*n* = 4) mouse brains were dried of excess PBS and placed in 10 ml Falcon tubes. Tubes were filled with Fluorinert (FC-3283, 3M Electronics) for susceptibility matching and to improve shimming.

Data were acquired at 9.4 T (20 cm internal diameter, horizontal bore, Bruker BioSpec Small Animal MR System, Bruker Biospin, Billerica, MA) using a 6 cm high performance gradient insert (maximum gradient strength: 1000 mT/m, Bruker Biospin) and a 35 mm internal diameter quadrature volume coil (Rapid MR International, Columbus, Ohio). Third-order shimming was iteratively performed over an ellipse that encompassed the entire brain using the Paravision mapshim protocol. *B*_0_ maps were produced by recording the voxel-wise frequency of the peak of the resonance, including additional sub-spectral resolution frequency produced by estimating the maximum peak amplitude of the resonance, described below.

Diffusion MRI (dMRI) was performed using a conventional 3D spin-echo/Stejskal-Tanner diffusion-weighted sequence (TR = 600 ms, TE = 11.389 ms, b-value = 3000 s/mm^2^, *δ* = 3.09 ms, Δ = 6 ms, spatial resolution = 125 µm isotropic, number of b0s = 8, number of directions = 30, receiver bandwidth = 150 kHz, duration = 36h 28min 48s).

3D-EPSI data were acquired using a MGE sequence with an oscillating readout gradient train. Sequence parameters were chosen so that the entire voxel-wise free induction decay was sampled to the noise floor with sufficiently high temporal resolution to ensure a large spectral bandwidth of ± 360 Hz around the main water peak (TR = 1000 ms, TE of first echo = 2.74 ms, echo spacing = 2.74 ms, number of echoes = 192, receiver bandwidth = 75 kHz, flip angle = 68°, 100 µm isotropic resolution, four averages, duration = 12 h). The average signal to noise ratio (SNR) is shown as a function of TE in Supporting Information Figure S1.

### 2.3 EPSI data processing

The following EPSI data processing and data analysis steps have also been reported elsewhere (15, 16) but are summarized here for completeness. Modifications reflecting differences in this work have been made.

EPSI data were processed and analyzed with IDL (ITT Visual Information Solutions, Boulder CO), Matlab (The MathWorks Inc., Natick, MA, 2012), and FSL (FMRIB Software Library, FMRIB, Oxford, UK). Data were processed to produce voxel-wise water spectra. Each 4D complex array (*k*_*x*_ × *k*_*y*_ × *k*_*z*_ × *t*) was Fourier transformed in all dimensions to produce three spatial dimensions and one spectral dimension (*x* × *y* × *z* × *ν*). For the complex spectrum, *ν*, spectral ghosting was corrected at each point in space (x, y, z) (43). The maximum peak magnitude was estimated in each voxel by applying the Fourier shift theorem to the complex data; the addition of a linear phase term in the temporal domain performs sub-spectral resolution shifts in the frequency domain allowing for the identification of the maximum signal magnitude for voxels in which the peak was located between Fourier components (44). This process was alternately iterated with a zeroth-order phase correction to produce pure absorption spectra (45).

Water peak height (PH) images were constructed with image contrast produced by the maximum voxel-wise signal amplitude of the water spectrum (46). This can be achieved by shifting the position of the maximum peak amplitude of the water resonance to the central Fourier component of the frequency axis. This step also serves to eliminate any relative background field information from each spectrum with little computational effort; this is analogous to implementing a background field removal technique, such as the PDF (29) or the SHARP (47) filter, to 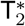-weighted gradient echo data processed in the temporal domain.

The EPSI datasets contained a number of regions with signal drop-out resulting from the magnetic susceptibility mismatch between the tissue and bubbles stuck to the surface of the brain or trapped in the ventricles. To exclude these regions from downstream analysis, artifact masks were constructed for each dataset using the `Atropos` tissue segmentation algorithm in the ANTs (48) software package on the water peak-height images. First, brain masks were automatically generated using the `bet` protocol in FSL (49). The algorithm was then initialized with a three-class K-means classification of the water peak-height images, with the classes representing white matter, gray matter, and joint CSF/artifact. Voxels in the CSF/artifact class were excluded from all further analysis. A sample image showing tissue classification and artifact filtering is shown in Supporting Information Figure S2.

### 2.4 Model fitting

Myelin imaging with MGE data has typically relied on the use of a three-compartment model for white matter, with separate 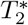 and frequency shifts stemming from axonal water, extracellular water, and myelin water. At its simplest, the time-dependent magnitude |*S*_*i*_(*t*)| of the MGE signal at voxel *i* is modeled as a sum of exponentials for each compartment (9, 17):

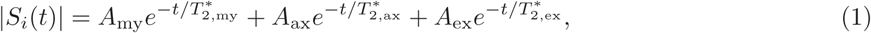

where *A*_my_, *A*_ax_, and *A*_ex_ refer to the amplitudes of the myelin, axonal, and extracellular compartments in voxel *i*, respectively.

In Van Gelderen et al. (31), the model was modified to include two frequency offsets for myelin (Δ*f*_my−ex_) and axonal (Δ*f*_ax−ex_) water relative to extracellular water. This model was also fit to magnitude data:

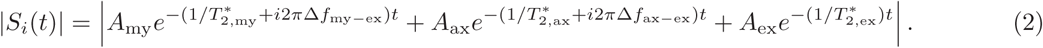

In Nam et al. (12), the model was fit to the full complex data and further extended to include frequency offset terms for all three compartments with respect to the background as well as a background phase term *ϕ*_0_:

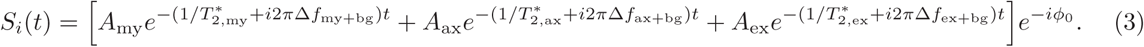

Our analysis focused on these final two models, which for simplicity we will refer to as the “magnitude fit” (Eqn. 2) and “complex fit” (Eqn. 3). Note that though Eqn. 2 is fit to magnitude data, it still represents a complex model with frequency shifts that produce asymmetric broadening of the water spectrum.

Model fitting was performed by first converting the preprocessed EPSI spectral data into the temporal domain using an inverse fast Fourier transform. The two models were then fit to the resulting voxel-wise FIDs in Python using a non-linear least-squares approach implemented with the `curve_fit` function in the SciPy package. Optimization parameters for the fitting were identical to those presented in Table 1 of Nam et al. (12) and are available for reference in Supporting Information Table S1.

### 2.5 Asymmetry

To quantify asymmetric broadening of the water resonance, we used a unitless spectral asymmetry metric (15, 16, 50, 51). At each voxel, the high-field half of the spectrum is subtracted from the low-field half and normalized by the total integral of the spectrum.

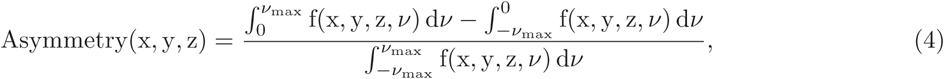

where *f* (*x, y, z, ν*) is the value of the water spectrum at a given position (*x, y, z*) and frequency *ν*. Integration was performed to *ν*_max_ = ±38 Hz from the main water peak (identified at 0 Hz for simplicity) to ensure that resonance details are captured while still reaching the spectral baseline. This cutoff value was shown in a previous study (16) to lead to asymmetry values sensitive to the differences between control and shiverer white matter. Overall, asymmetry results are robust to the specific choice of threshold (Supporting Information Figure S3). Before calculating asymmetry from the biophysical models, spectra were first computed by evaluating the analytic time-domain models at the echo points measured in the data using the parameters estimated for each voxel, then taking a fast Fourier transform.

### 2.6 Myelin water fraction

Three-compartment models are commonly fit to MGE data in order to calculate the myelin water fraction (MWF) metric, taken as the ratio of the myelin-compartment amplitude *A*_my_ to the total sum of amplitudes from each compartment:

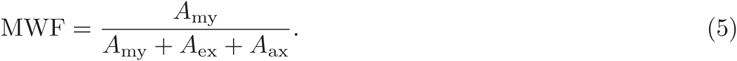

This metric was calculated at each voxel from the amplitude parameters estimated from both the magnitude- and complex-fit models in order to contextualize the utility of the asymmetric metric.

### 2.7 dMRI processing

#### 2.7.1 fODF fitting and registration

dMRI processing was performed with the MRtrix3 software package (52). Data were denoised using the `dwidenoise` routine (53, 54). Binary brain masks were generated with the `dwi2mask` routine to speed further processing. The datasets were fit to a tensor model (55) using `dwi2tensor` to calculate the fractional anisotropy (FA) metric used as a proxy for white matter content. The data were then fit to fiber orientation distribution functions (fODFs) using constrained spherical deconvolution (56, 57) to estimate the orientation of the principal fiber populations and to evaluate the presence of additional crossing fiber populations within each voxel. Comparison of fODFs across datasets requires global intensity normalization of the diffusion data prior to reconstruction, since the data are not first log-normalized with the *b*_0_ volume. First, the data were bias-corrected using the `N4BiasFieldCorrection` algorithm (58) in ANTs, then global intensity normalization was done with the `dwinormalise group` routine in MRTrix3. White matter response functions were then calculated for each dataset using the `tournier` algorithm (59), with *ℓ*_max_ = 6 (28 coefficients). The group-averaged response function was then used to fit the bias-corrected diffusion-weighted images to fODFs with the `dwi2fod` command.

The FA images were spatially registered to the corresponding EPSI peak-height images using affine transformations calculated in ANTs using a mutual information maximization approach. The FA images were chosen for registration because they exhibit greater white/gray matter contrast, particularly for the shiverer data. The resulting affine transformations were used to warp, reorient (60), and modulate (61) the fODFs using the `mrtransform` command in MRTrix3, which preserves apparent fiber densities across fODF lobes before and after spatial transformation.

#### 2.7.2 Microstructural analysis

As in Foxley et al. (16), voxels across the entire dataset were binned according to FA value with thresholds of FA ≤ 0.3, 0.3 < FA ≤ 0.45, 0.45 < FA ≤ 0.6, and FA > 0.6. Visual inspection of the resulting voxel masks suggests that the lower FA ≤ 0.3 bin consists of predominately gray matter while the upper FA > 0.6 bin consists of predominately white matter, with the additional two bins composed of mixed populations. To account for known biases in FA in voxels with crossing fibers (62), voxels were further characterized by the number of distinct fiber populations. Individual lobes of each fODF were segmented in MRTrix3 using the `fod2fixel` and `fixel2peaks` commands. The number of populations (*N*_*f*_) at each voxel was recorded as either “single” (*N*_*f*_ = 1) or “crossing” (*N*_*f*_ > 1). To explore the effect of susceptibility anisotropy (22– 26) in the model-derived spectra, the angle between principal fiber orientations and *B*_0_ was calculated as Γ = cos^−1^ (*s*_*z*_), where ***ŝ*** = (*s*_*x*_, *s*_*y*_, *s*_*z*_) is the fiber orientation unit vector at a given voxel and *B*_0_ points along 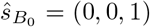. Voxels with both single and crossing fibers were pooled by angle into bins in 5° increments based on the orientation of the primary fiber population. The voxel counts across all values of FA, *N*_*f*_, and Γ are shown for both control and shiverer data in Supporting Information Figure S4.

### 2.8 Statistical analysis

We evaluated the sensitivity of the data- and model-derived spectra to the differences between control and shiverer white matter by considering asymmetry as a one-variable binary classifier for control or shiverer data and using the area (AUC) under the receiver operating curve (ROC) as a function of FA and the number of fiber populations. For this analysis, ROC curves were created by taking the distribution of voxel-wise control and shiverer asymmetries for each FA bin and fiber population number group, varying the asymmetry threshold value used to identify test voxels from the control datasets, and calculating the overall sensitivity and specificity at each threshold. The AUC was then calculated by numerical integration of the ROC curve. This procedure was repeated for the MWF metric for comparison to asymmetry.

## 3 Results

### 3.1 Comparison of asymmetry values

Figure 1 shows the distribution of asymmetry values calculated from the raw data and both models across all voxels from control and shiverer samples. At the whole-brain level there is a small but clear separation between control and shiverer asymmetry values observed in the data, whereas both model-derived asymmetry distributions are virtually indistinguishable between control and shiverer. Spectra derived from each of the two models also dramatically underestimate the range of asymmetry magnitudes observed in the data. Across all voxels in both tissue types, data-derived asymmetry has a standard deviation of 0.0427, while the magnitude- and complex-fit asymmetries have standard deviations of 0.0140 and 0.0058, respectively.

**Figure 1:**
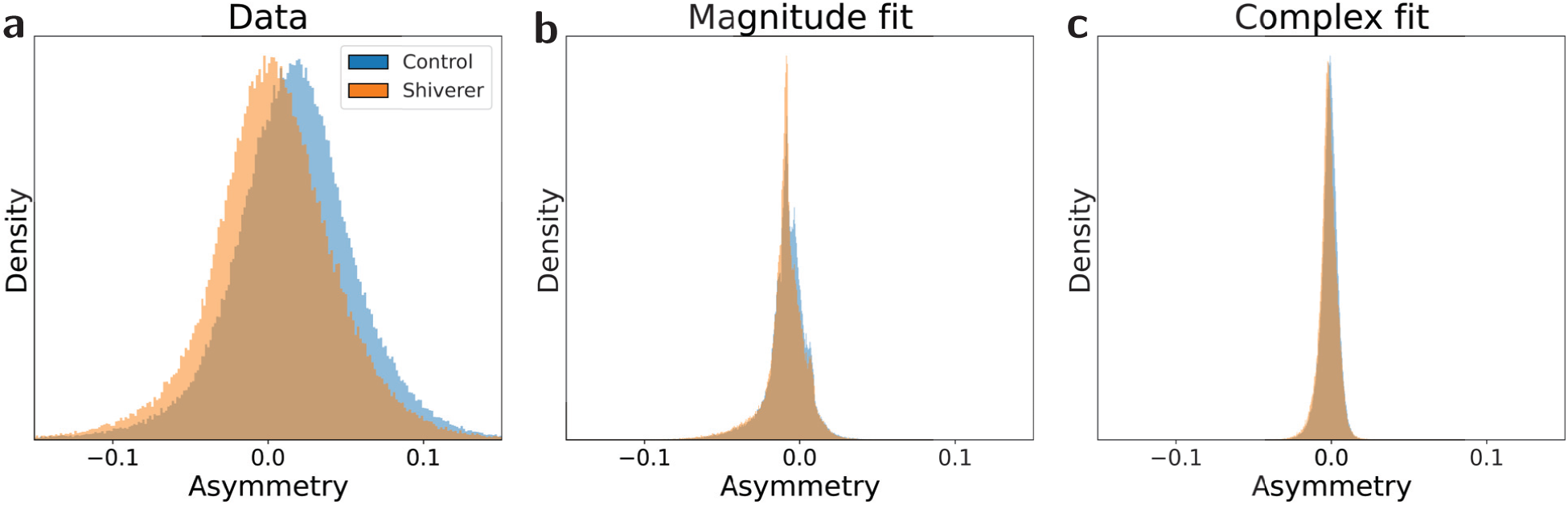
Histograms of asymmetries from spectra derived from (a) data, (b) the magnitude-fit model, and (c) the complex-fit model.

Voxel-wise comparisons between data- and model-derived asymmetries are shown as scatterplots across different FA bins in Figure 2. Asymmetries from the magnitude-fit model show negligible correlation (R^2^ ≈ 0) with data-derived asymmetries from both control and shiverer samples across all FA bins. The complex-fit model agrees slightly better with the data, with R^2^ values that increase with increasing FA for control data up to a maximum of R^2^ = 0.403 for the highest FA bin, while complex-fit R^2^ values for shiverer data are lower than for control data and do not increase with FA. All R^2^ values were found to be significant at the *α* = 0.1 level using an F-test.

**Figure 2:**
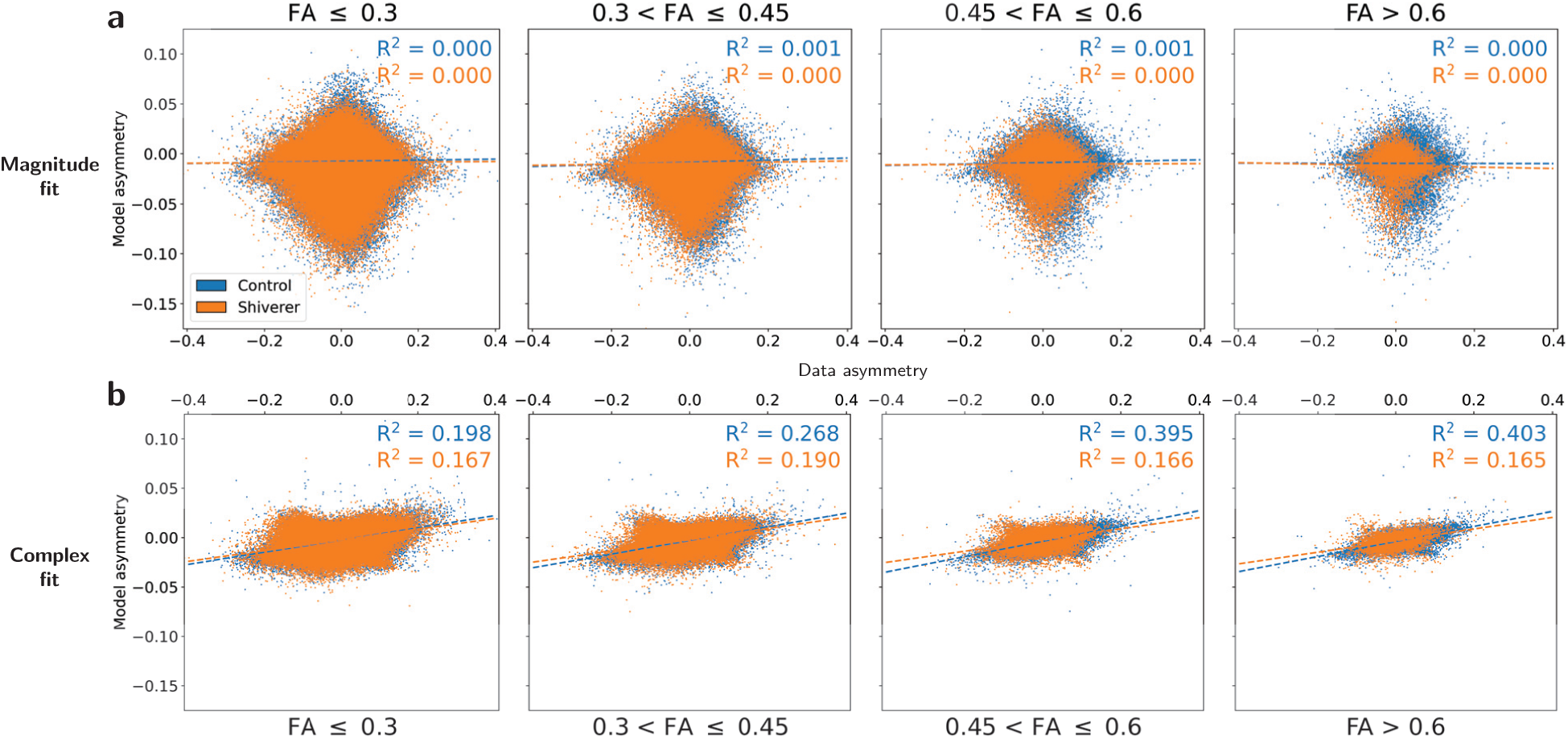
Scatterplots of data- and model-derived asymmetries for the (a) magnitude-fit and (b) complex-fit models. Dashed lines represent the results of linear regressions and the R^2^ values are the square of the Pearson correlation coefficients.

Figures 3a–b show 2D histograms of the absolute asymmetry difference between the models and data vs. the adjusted R^2^ assessing the goodness-of-fit of the models to the FID data for voxels with FA > 0.6. Both models generally fit the FID data very well overall, with the mean adjusted R^2^ values across all voxels from both tissue types being 0.994 and 0.988 for the magnitude-fit and complex-fit models, respectively. However, the high adjusted R^2^ of the models in the temporal domain does not correspond to accuracy in reproducing the asymmetric broadening observed in the spectral domain of the data — correlations between the adjusted R^2^ and asymmetry difference are negligible for both models. This effect is visually demonstrated further in Figure 3c, which shows two representative white-matter voxels from the anterior commissure tract of a control mouse. In both sample voxels, the two models fit very closely to the data in the temporal domain (adjusted R^2^ > 0.98) but greatly underestimate asymmetry in the frequency domain. For example, in the case of the voxel highlighted in green, both models clearly miss a prominent secondary peak around 8 Hz (black arrow in Figure 3c, bottom right) corresponding to an oscillation in the FID around 0.05-0.18 seconds.

**Figure 3:**
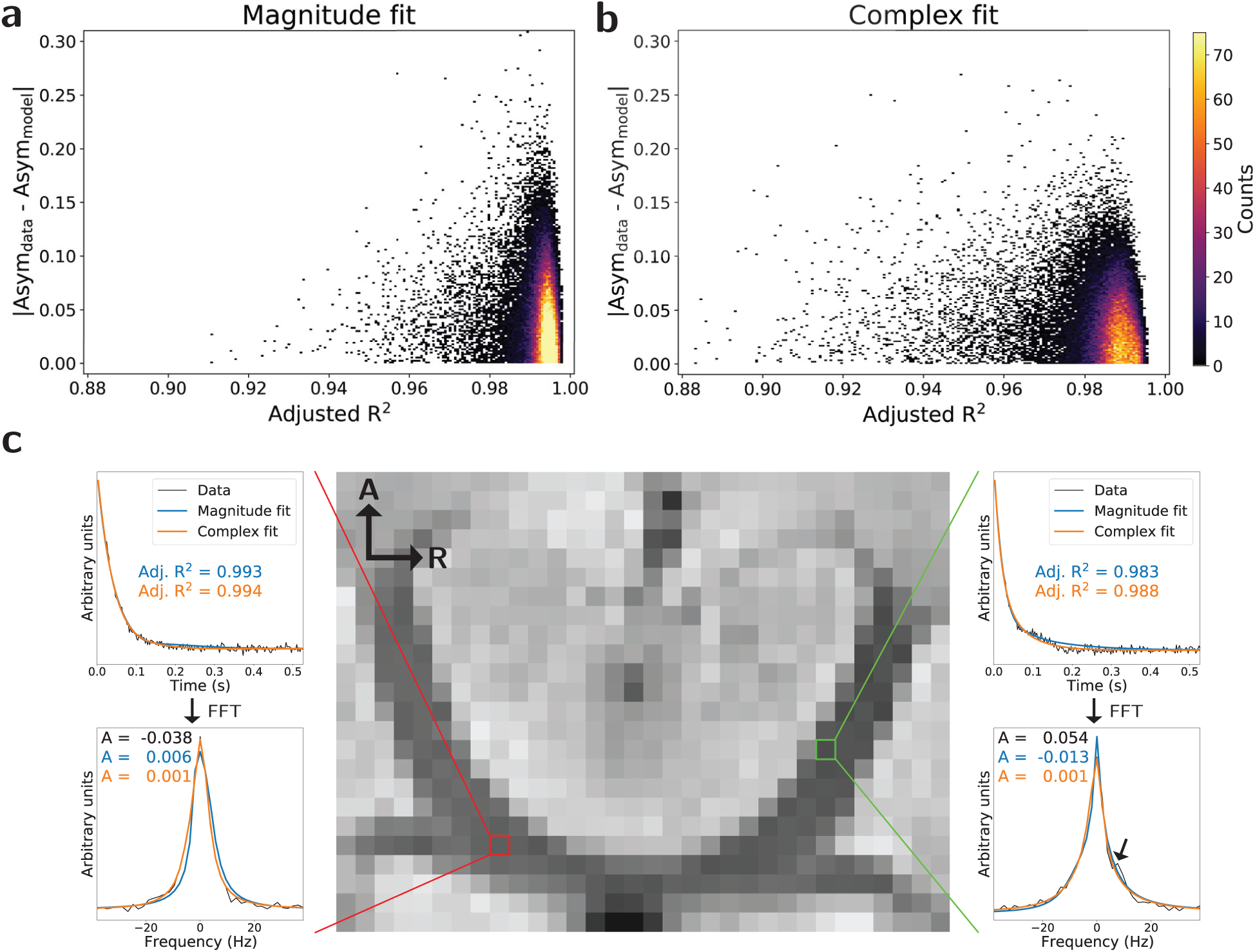
(a-b) 2D histograms of the absolute difference between measured and model-estimated asymmetries vs. the model goodness-of-fit in terms of adjusted R^2^ for the (a) magnitude-fit and (b) complex-fit models for voxels with FA > 0.6. Green dashed lines show linear regressions with associated *R*^2^ values indicating a negligible relationship. (c) Water peak-height image showing the anterior commissure tract of a control mouse with two representative voxels demonstrating the mismatch between the model goodness-of-fit and asymmetry difference. A–R axis labels correspond to the Anterior and Right directions, respectively.

Model fits were further evaluated with the Bayesian information criterion (BIC). The distributions of the differences in BIC between the magnitude-fit model (8 free parameters) and the complex-fit model (10 free parameters) are shown in Supporting Information Figure S5. For all FA bins and both control and shiverer data, BICs were lower from the magnitude-fit model than the complex-fit model.

Figure 4 shows additional 2D histograms for high FA (FA > 0.6) voxels demonstrating the relationship between the asymmetry directly measured in the spectral data and the compartmental frequency shifts predicted by the models. Note that the magnitude-fit model (Eqn. 2) includes frequency shift terms for two compartments: Δ*f*_my−ex_ and Δ*f*_ax−ex_, defined as the difference between the myelin and axonal water shift with the the extracellular water shift, respectively. The complex-fit model (Eqn. 3) includes frequency shift terms for all three components: Δ*f*_my+bg_, Δ*f*_ax+bg_, Δ*f*_ex+bg_, defined as the additional shifts above a background. For consistent comparison, the extracellular shifts have been subtracted from the complex-fit model frequencies reported in Figure 4. The magnitude-fit frequencies again show virtually no correlation with the measured asymmetry, while the complex-model frequencies show only weak correlation (R^2^ = 0.114) in the axonal compartment.

**Figure 4:**
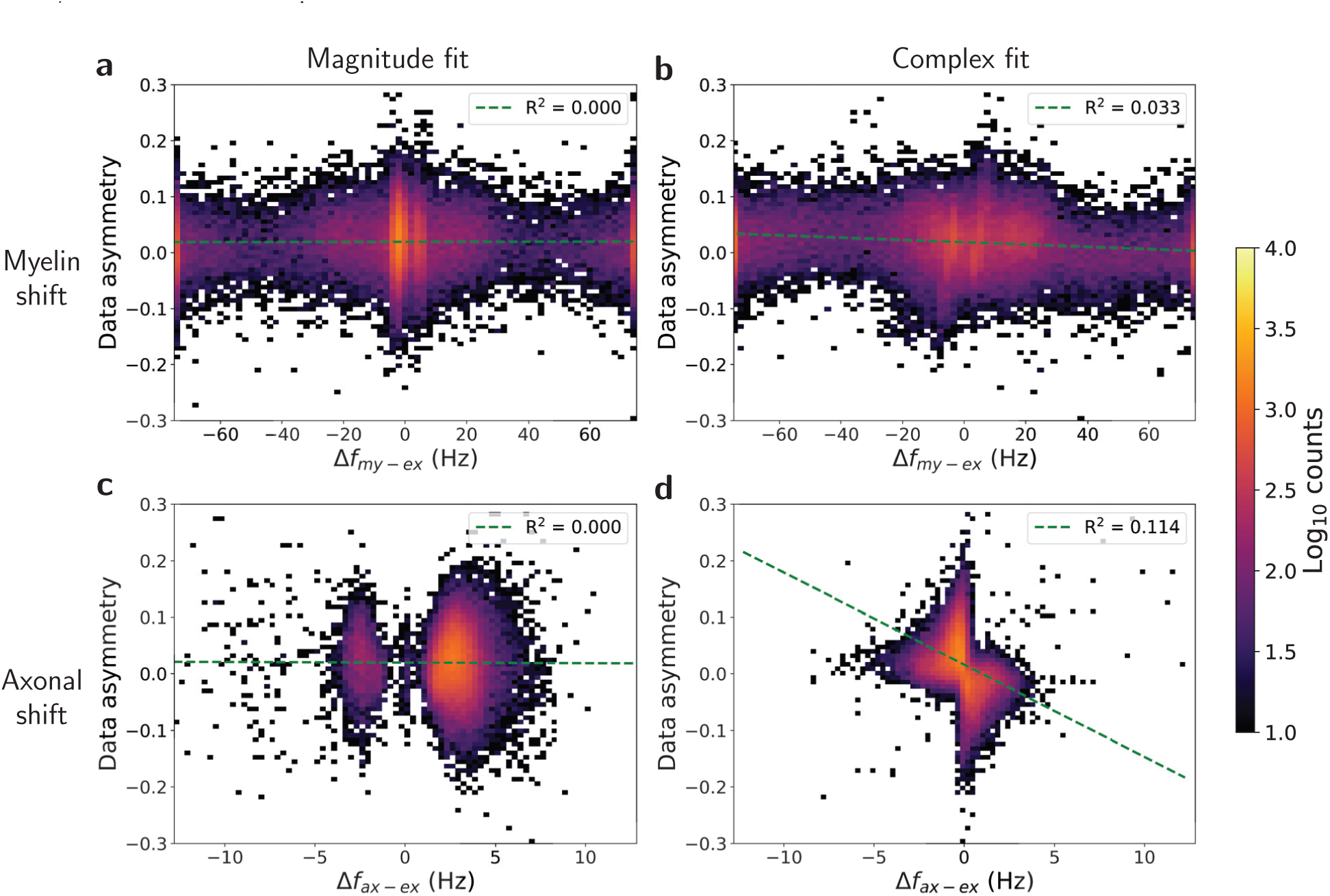
2D histograms showing the relationship between data asymmetry and model-predicted frequency shifts for the myelin and axonal water compartments in voxels with FA > 0.6. Green dashed lines show linear regressions with associated R^2^ values.

### 3.2 Sensitivity to shiverer white matter

While Figure 1 showed the range of asymmetry values across the entire dataset, Figure 5 shows asymmetry distributions grouped by FA bin for single- and crossing-fiber voxels. Previous work has shown that data-derived spectral asymmetry is sensitive to white-matter differences between control and shiverer mice (16), with spectra exhibiting consistent upfield broadening along white-matter tracts in control mice that decreases in magnitude in shiverer mice. This is demonstrated in Figures 5a–b, which show an increasing separation between control and shiverer data-derived asymmetries as FA increases, independent of the number of fiber populations. This effect is not observed to the same extent after fitting the data to models, as shown in Figures 5c–f. The separation between the control and shiverer distributions is quantified in Figure 6 with AUCs resulting from ROC analysis. While separation between control and shiverer distributions did increase marginally with FA for both models, data-derived asymmetry led to a substantially higher AUC than either of the model-based approaches under all microstructural conditions. The classification performance of data-derived asymmetry was robust to the number of fiber populations, with only slight decreases in AUC for higher FA voxels with crossing fibers (FA > 0.45) compared to single fibers and slight increases for lower FA voxels with crossing fibers (FA ≤ 0.45) compared to single fibers, potentially owing to the fact that voxels with tightly packed, coherent fibers are likely to lead to artificially low FA values if they contain multiple distinct populations.

**Figure 5:**
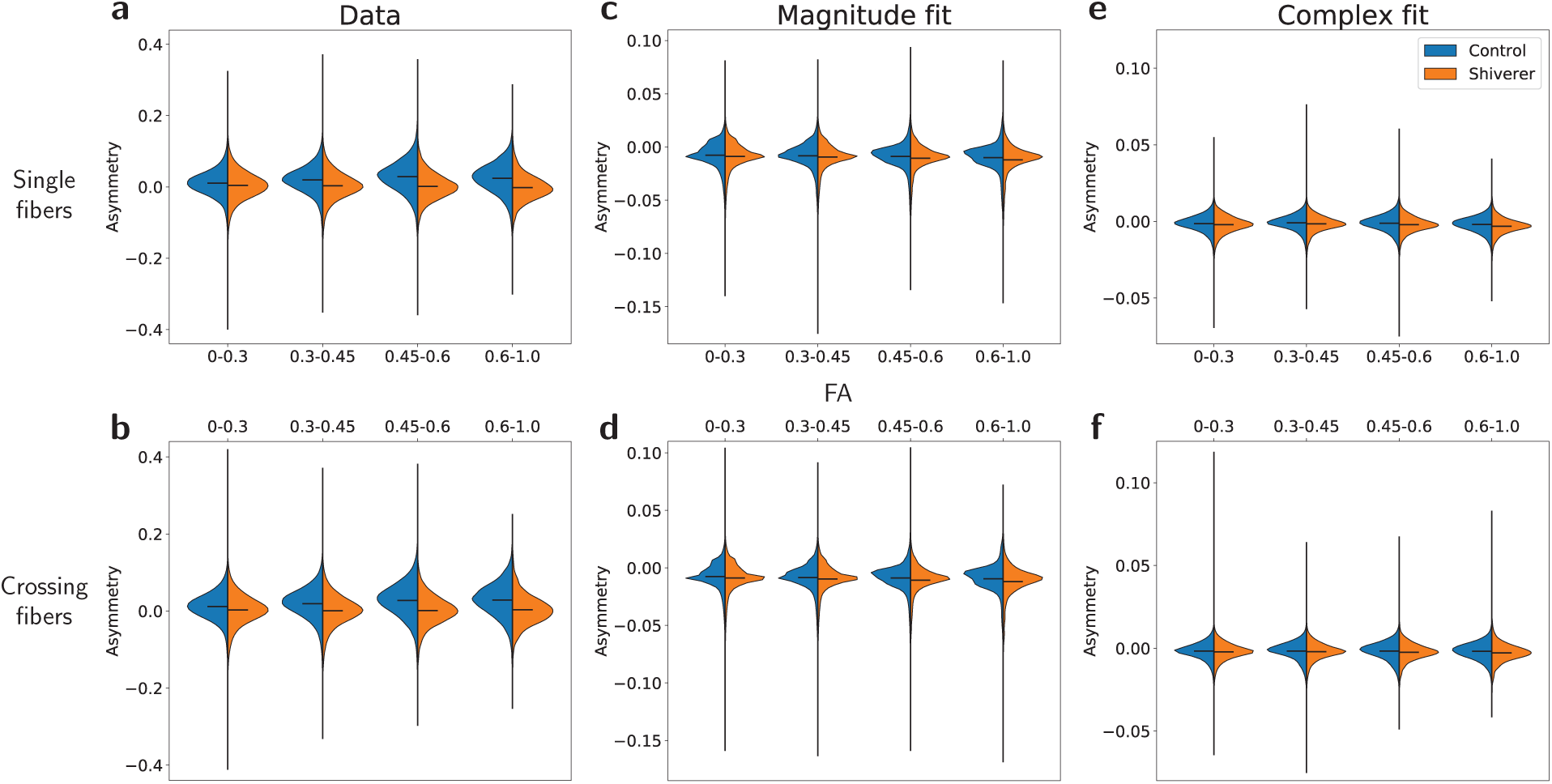
Violin plots illustrating distributions of control and shiverer asymmetries derived from (a–b) data, (c–d) the magnitude-fit model, and (e–f) the complex-fit model as a function of FA bin for voxels with single (a,c,e) and crossing (b,d,f) fibers. Distribution means are indicated by black lines. Note the difference in y-axis limits between (a–b), (c–d), and (e–f).

**Figure 6:**
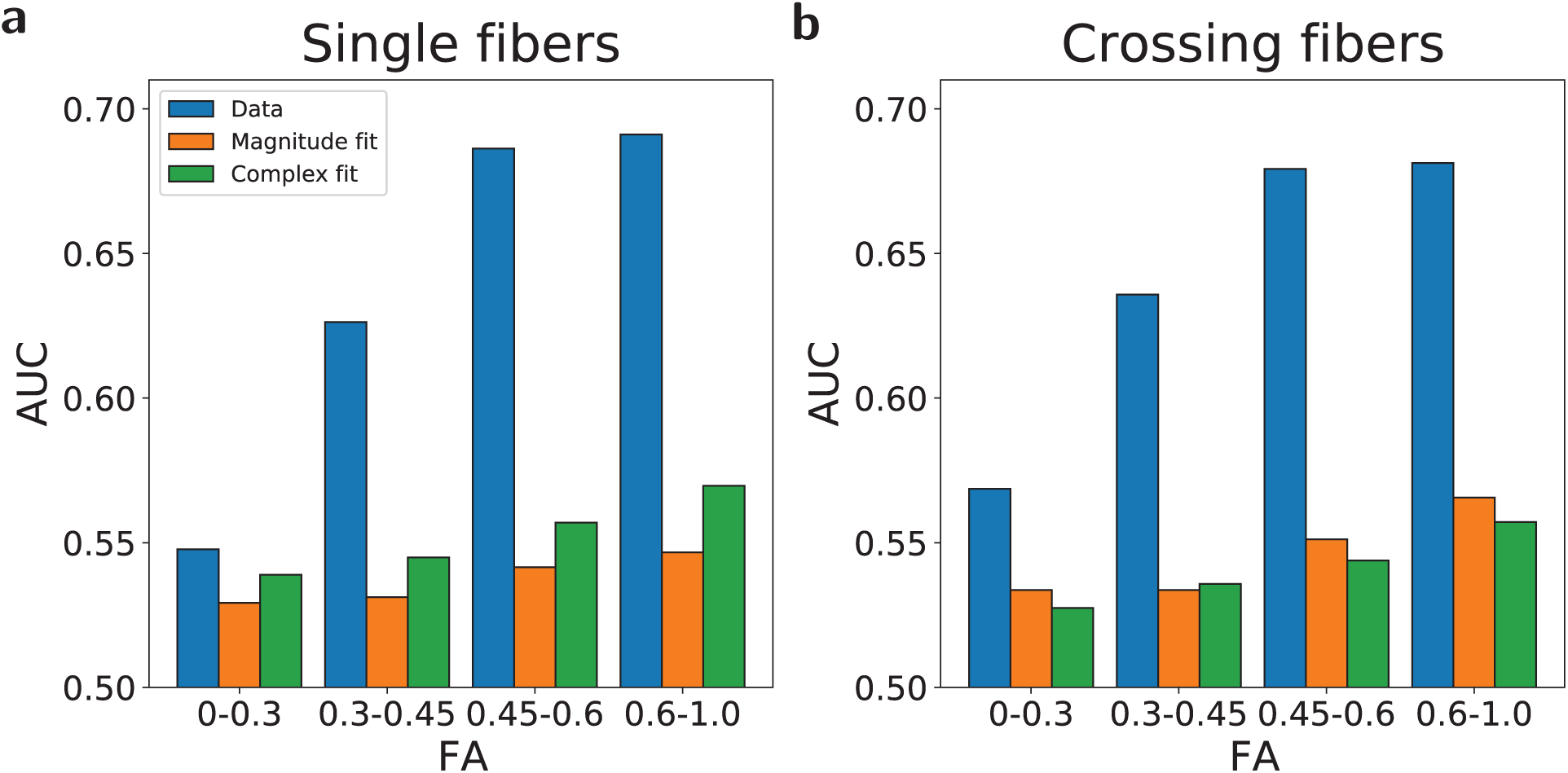
Values for the area under the ROC curve (AUC) using asymmetry as a one-variable classifier for control vs. shiverer data. Values represent AUCs for subsets of voxels in different FA bins containing either (a) single or (b) crossing fiber populations.

In Figure 7, data-derived asymmetry AUCs are compared to AUCs resulting from ROC analysis performed using the MWF metric from both models. Data asymmetry led to a greater separation between control and shiverer data than both model-derived MWF values for low-FA voxels (FA ≤ 0.6), while asymmetry and magnitude-fit MWF AUCs were comparable for high-FA voxels (FA > 0.6).

**Figure 7:**
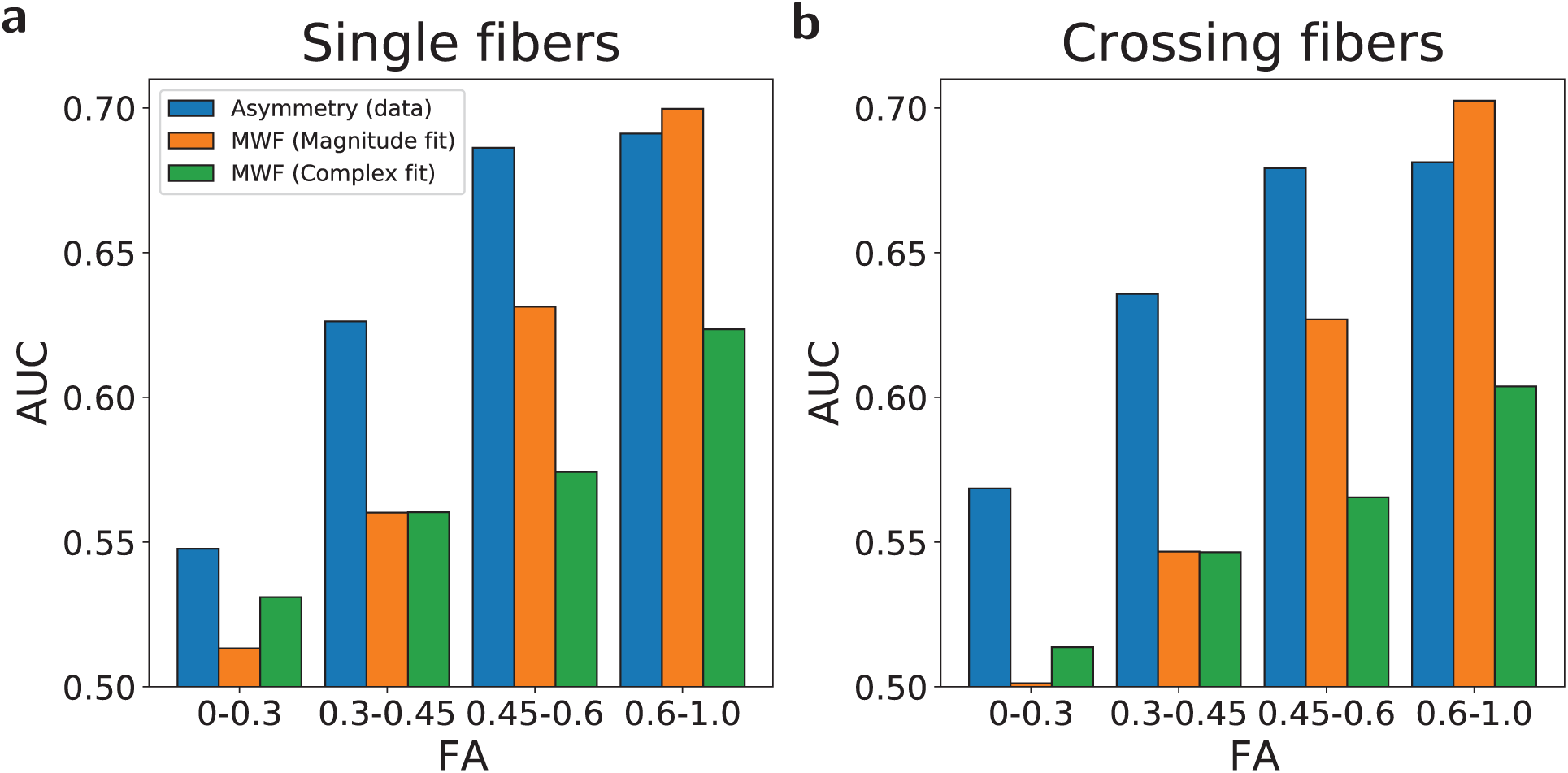
Values for the area under the ROC curve (AUC) using data asymmetry (blue) and model-based MWF (orange and green) as one-variable classifiers for control vs. shiverer data. Values represent AUCs for subsets of voxels in different FA bins containing either (a) single or (b) crossing fiber populations.

The white-matter sensitivity of the data-derived asymmetry is further demonstrated in the left column of Figure 8, which shows representative coronal slices of asymmetry images. While the data-derived asymmetry leads to observable gray/white-matter contrast in the control image comparable to the MWF images in Figures 8g–j, asymmetry images calculated from each of the models (Figures 8c–f) show markedly lower contrast, with virtually no distinguishable white matter tracts visible in the images derived from the magnitude-fit model in particular.

**Figure 8:**
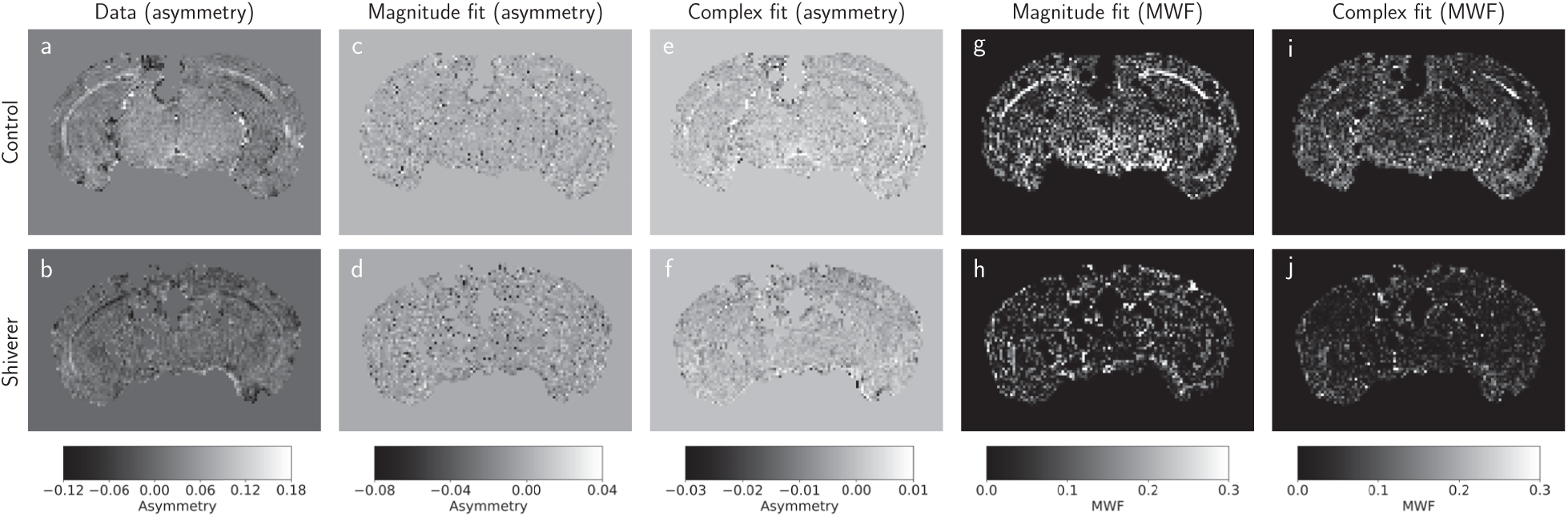
Representative coronal slices of images derived from (top) control and (bottom) shiverer samples. Images represent spatial distributions of (a–b) data-derived asymmetry, (c–d) magnitude-fit asymmetry, (e–f) complex-fit asymmetry, (g–h) magnitude-fit MWF, and (i–j) complex-fit MWF.

To explore the orientation-dependence of the model-estimated spectra, Figure 9 shows the mean asymmetry as a function of Γ, the angle between the orientation of the primary fiber population and *B*_0_ for voxels with FA *≥* 0.6. Color-shaded regions represent the interquartile range (25–75th percentile) across all voxels within each angular bin, and gray-shaded regions represent regions where the difference between control and shiverer values was not found to be statistically significant using a t-test with *α* = 0.01 after correcting for multiple comparisons. Both the data and the complex-fit asymmetry values show a clear relationship between asymmetry and Γ with good agreement with the susceptibility anisotropy model (24),

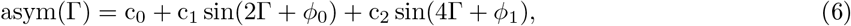

though the angular effect is much less pronounced for the magnitude-fit model and both models once again show far less separation between control and shiverer values than observed from the data before model-fitting. Note that data-based adjusted R^2^ values to the susceptibility anisotropy model are slightly lower for voxels with crossing fibers (Figure 9b) than for voxels with single fibers (Figure 9a), again suggesting that the performance of macroscopic imaging models are sensitive to the microstructural geometry of the underlying tissue.

**Figure 9:**
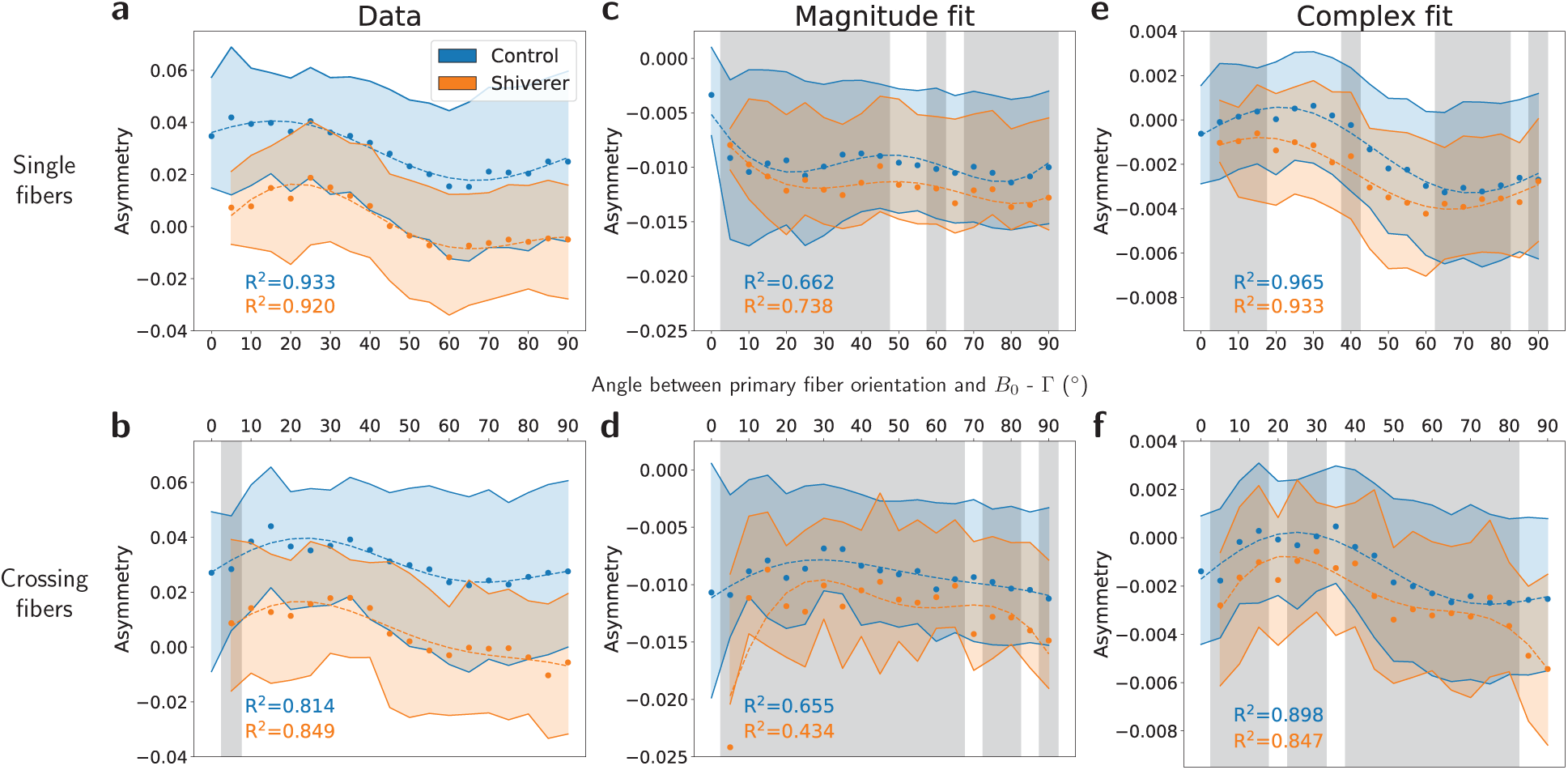
Relationship between asymmetry and Γ, the angle between the orientation of the primary fiber population and *B*_0_ for (a–b) data (c–d) the magnitude-fit model, and (e–f) the complex-fit model. Points represent averages within angular bins with a width of 5° across all voxels with FA *≥* 0.6 containing either (a,c,e) single fibers or (b,d,f) crossing fibers. Color-shaded regions represent the interquartile range (25-75th percentiles) across all voxels. Dotted lines represent fits to the susceptibility anisotropy model with associated adjusted R^2^ values. Gray regions represent angular bins for which the difference between control and shiverer values was not found to be significant with a t-test at *α* = 0.01.

## 4 Discussion

This study aimed to characterize the extent to which water spectra derived from two biophysical signal models fit to EPSI data are able to replicate spectral characteristics observed directly from the data itself, specifically with respect to sensitivity to white-matter differences between control and shiverer mice. We used spectral asymmetry as a summary metric to explore how data- and model-derived spectra differ over a range of microstructural environments. Our overall finding is that independent of how well the models fit the temporal data, the spectral asymmetry predicted by both models showed measurable discrepancies with the values observed in the data. The simplicity of the models provides interpretability and computational advantage at the expense of important disagreements with the data in the spectral domain. Specifically, the models dramatically underestimated the magnitude of the asymmetric broadening effect (Figure 1), with model-derived values for asymmetry that only loosely correlate with those measured in the data in high-FA voxels (Figure 2). Most importantly, the process of fitting the data to these simple biophysical models effectively leads to compromised spectral sensitivity to changes in white matter structure under all microstructural conditions explored in this work (Figures 6 and 8).

While the above conclusions apply broadly to both models, the complex-fit model did agree slightly better with the data with respect to certain spectral characteristics, a finding consistent with previous work comparing these two models (12). Complex-fit asymmetries were correlated more strongly with data-measured asymmetries across all FA bins (Figure 2) than magnitude-fit asymmetries were. Complex-fit asymmetry correlations increased slightly with FA in control data but not in shiverer data, indicating the expected result that the model agrees best with the data under white-matter microstructural conditions most similar to the assumed biophysical model, under the assumption that high-FA control voxels are likely to contain myelinated, coherent white-matter tracts. Similarly, the axonal water compartment frequency shift estimated from the complex-fit model was the only compartmental frequency shown to have any correlation with the raw-data asymmetry (Figure 4). Complex-fit asymmetries were also shown to have a stronger relationship to the angle between the fiber orientation and *B*_0_ than magnitude-fit asymmetries (Figure 9). Overall, however, both models showed relatively poor performance with respect to spectral white-matter sensitivity and contrast (Figures 6 and 8). The complex-fit model also showed slightly worse performance than the magnitude-fit model with respect to white-matter sensitivity using the MWF metric, as well as consistently lower values for the Bayesian information criterion (BIC) assessing model fit (Supporting Information Figure S5).

The data-derived asymmetry results presented in this work are consistent with previous studies demonstrating the utility of model-free analysis of fully-sampled MGE/EPSI data in the frequency domain towards white-matter imaging (14–16). Our results indicate that biases are present in the explored compartmental models even at the level of compressing the full spectral information into a single scalar asymmetry metric, promoting caution in the downstream interpretation of model-predicted spectra.

Notably, this work also demonstrates the robustness of the asymmetry metric under complex microstructural conditions. Measured asymmetries were shown to have comparable sensitivities to white matter (Figure 6) and demonstrate similar behavior with respect to fiber angle (Figure 9) in voxels with single and crossing fibers. A direction of future work will be to more rigorously characterize the spectral response to the specific number as well as relative strength and position of fiber populations within the voxel.

As is common with multi-exponential fitting with MGE data, SNR is a limitation when fitting of models with so many parameters. The discrepancies between models in our results promote the more widespread use of direct validation of model parameters using simulation studies and histological ground truth imaging. For this work, this validation role was played by the known physical differences between control and shiverer mice and the empirical sensitivity to those differences present in the data prior to model fitting. We also hypothesize that the improvement of future models will also rely on more advanced noise modeling and optimization protocols to account for the non-Gaussian distribution of the data noise, particularly for the magnitude model. In this work, our approach was instead to fairly benchmark both models by applying them as they were originally published.

This analysis method of benchmarking spectroscopic MR data in the temporal domain against data in the spectral domain can be extended to existing datasets after applying a linear transform to the FID to produce absorption spectra. We note, however, the importance of sampling to the results in this work – the sensitivity of the spectral analysis is dependent on the increased spectral resolution that comes by sampling the FID into the noise floor, which also helps to avoid time-domain truncation effects which could potentially produce ringing artifacts in the spectra that could be mistaken for spectral asymmetry. Supporting Information Figure S6 shows AUCs similar to Figure 6, and demonstrates that after truncating the FID to just the first 32 echoes, neither the data-nor the model-derived asymmetries show meaningful sensitivity to shiverer white matter, while Supporting Information Figure S7 shows that the AUCs stabilize for data-derived asymmetry around 64 echoes.

While we are unable to isolate myelin as a specific driver of spectral asymmetry without independent histological validation of myelin content, we hypothesize that differences in myelin content and structure between control and shiverer mice do in fact play a sizable role both in this effect and in additional findings of this study. Multiple previous studies have also compared MR methods between control and shiverer specimens with histological validation (33–35). Specifically Takanashi et al. (35) performed immunohistochemical analysis of shiverer mice brains that revealed an almost total lack of myelin in the thalamus, white matter, and cortex. These results correlated with findings from MR spectroscopy data. Shiverer mice did demonstrate astrogliosis and an increased number of total oligodendrocytes in the white matter, though we do not expect these changes to affect the asymmetric broadening of the spectrum since there is nothing inherent to these cell types that would drive a local change in magnetic susceptibility (25, 63).

Overall, we feel that our analysis reveals that there is underutilized spectral information within EPSI data that yields important insight to white-matter structure complementary to information from traditional temporal-domain metrics such as MWF. This motivates the further development of spectral-domain components of biophysical white-matter models. Validation of models and physical parameter ranges have typically relied on simulations using a simple underlying geometric model of white matter that uses hollow cylinders to describe axons (13, 27, 28, 31). A recent study (18) simulated the MR signal arising from axons under different 2D geometric models. It showed that as the axon geometry model becomes more realistic through the use of segmented cross-sections from electron micrographs, the generated microscopic magnetic field becomes increasingly complex and the compartment-specific components of the water spectra become much less distinct, leading to asymmetric broadening around the main peak as opposed to narrow compartment-specific peaks. Though part of this effect could potentially be attributed to fixation-related warping of the axons imaged with electron microscopy (EM) for the study, these results suggest that microstructural imaging pipelines that rely on biophysical models might be biased by the models’ assumed geometries. Recent multi-modal analysis of mouse brain data (64) has demonstrated that the geometric complexity of axon populations increases dramatically in three dimensions; nerve fibers undulate, fan, and cross within MRI-sized-voxels. Such datasets will be important for future MR simulation studies that use realistic 3D white-matter models to explore the spectral signatures of tissue microstructure under different geometric configurations. In many results of this work, we observed that model performance was highly sensitive to the underlying tissue microstructure. The results of realistic 3D simulation studies could potentially guide the development of hybrid microstructural imaging approaches where structural information from dMRI data could be used to select relevant model components and parameters for EPSI. Some work towards this end is already underway, a recent study by Hédouin et al. (65) created a pipeline for creating realistic 2D white-matter models for MR simulation studies and a deep learning approach to recover microstructural properties with a combination of GRE and dMRI data. They validated their methods using a single 3D EM volume containing relatively coherent fibers from a mouse corpus callosum. Further validation against EM data representing more complicated geometries of crossing fibers could provide more robust insights into the role of microstructural geometry on the resulting MR signal.

## 5 Conclusion

Through analysis of fully-sampled MR spectra from control and dysmyelinated mouse brain, we reveal discrepancies between measured spectra and spectra estimated with existing biophysical compartmental models traditionally used for myelin imaging. We showed that spectra estimated from these biophysical models do not accurately predict the extent of asymmetric broadening observed in white-matter voxels, leading ultimately to compromised sensitivity to important differences in white-matter structure. This work further demonstrates the utility of model-free analysis of the water resonance spectrum with fully-sampled EPSI data, promoting its continued development as a tool to benchmark novel biophysical signal models.

## 6 Acknowledgments

S.T. is supported by the National Institutes of Health (NIH) (F31NS113571). N.K. is supported by the NIH Brain Initiative (U01MH109100). P.L.R. is partially supported by the NIH (R01EB026300). Additional funding was provided by the NIH (S10OD025081, S10RR021039, P30CA14599).

