## Supporting Information for "Model-free analysis in the spectral domain of postmortem mouse brain EPSI reveals inconsistencies with model-based analyses of the free induction decay"

Table S1: Replicated from Table 1 in Nam et al. (11). Initial values and search ranges of the parameters for the magnitude-fit and complex-fit models. $S_1 = S(\mathrm{TE}_1)$. $\Delta f_{\mathrm{bg,init}} = \angle \frac{\{\sum_{n=1}^{N-1} S_n^* S_{n+1}\}}{2\pi \Delta \mathrm{TE}}$: initial $\Delta f_{\mathrm{bg}}$ ($N$ = number of echoes used in fitting).

| | Both models | | | | | |
|---|---|---|---|---|---|---|
| | $A_{\mathrm{my}}$ | $A_{\mathrm{ax}}$ | $A_{\mathrm{ex}}$ | $T^*_{2,\mathrm{my}}$ (ms) | $T^*_{2,\mathrm{ax}}$ (ms) | $T^*_{2,\mathrm{ex}}$(ms) |
| Initial value | $0.1 \times \lvert S_1 \rvert$ | $0.6 \times \lvert S_1 \rvert$ | $0.3 \times \lvert S_1 \rvert$ | 10 | 64 | 48 |
| Lower bound | 0 | 0 | 0 | 3 | 25 | 25 |
| Upper bound | $2 \times \lvert S_1 \rvert$ | $2 \times \lvert S_1 \rvert$ | $2 \times \lvert S_1 \rvert$ | 25 | 150 | 150 |
| | Magnitude fit | | Complex fit | | | |
| | $\Delta f_{\mathrm{my-ex}}$ (Hz) | $\Delta f_{\mathrm{ax-ex}}$ (Hz) | $\Delta f_{\mathrm{my+bg}}$ (Hz) | $\Delta f_{\mathrm{ax+bg}}$ (Hz) | $\Delta f_{\mathrm{ex+bg}}$ (Hz) | $\phi_0$ (rad) |
| Initial value | 5 | 0 | $\Delta f_{\mathrm{bg,init}}$ | $\Delta f_{\mathrm{bg,init}}$ | $\Delta f_{\mathrm{bg,init}}$ | $\angle S_1$ |
| Lower bound | -75 | -25 | $\Delta f_{\mathrm{bg,init}} - 75$ | $\Delta f_{\mathrm{bg,init}} - 25$ | $\Delta f_{\mathrm{bg,init}} - 25$ | $-\pi$ |
| Upper bound | 75 | 25 | $\Delta f_{\mathrm{bg,init}} + 75$ | $\Delta f_{\mathrm{bg,init}} + 25$ | $\Delta f_{\mathrm{bg,init}} + 25$ | $\pi$ |

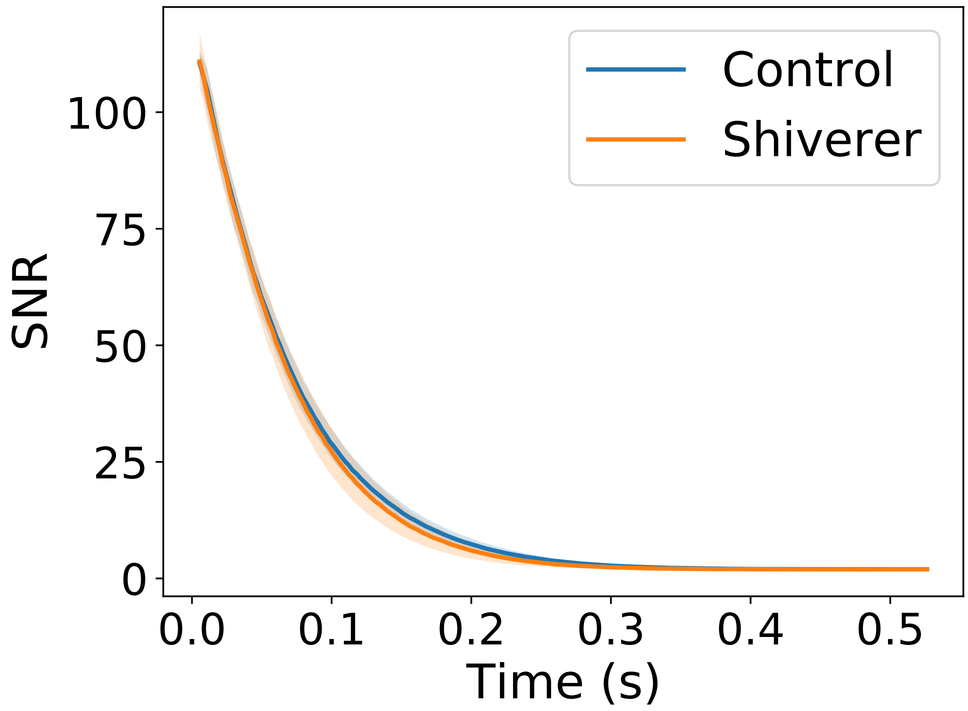


Figure S1: SNR as a function of echo time. SNR was measured by taking the ratio of the average in-brain signal volume and the standard deviation of a rectangular patch of background voxels at each TE. Lines represents averages across datasets and the shaded region represents ± 1 standard deviation.

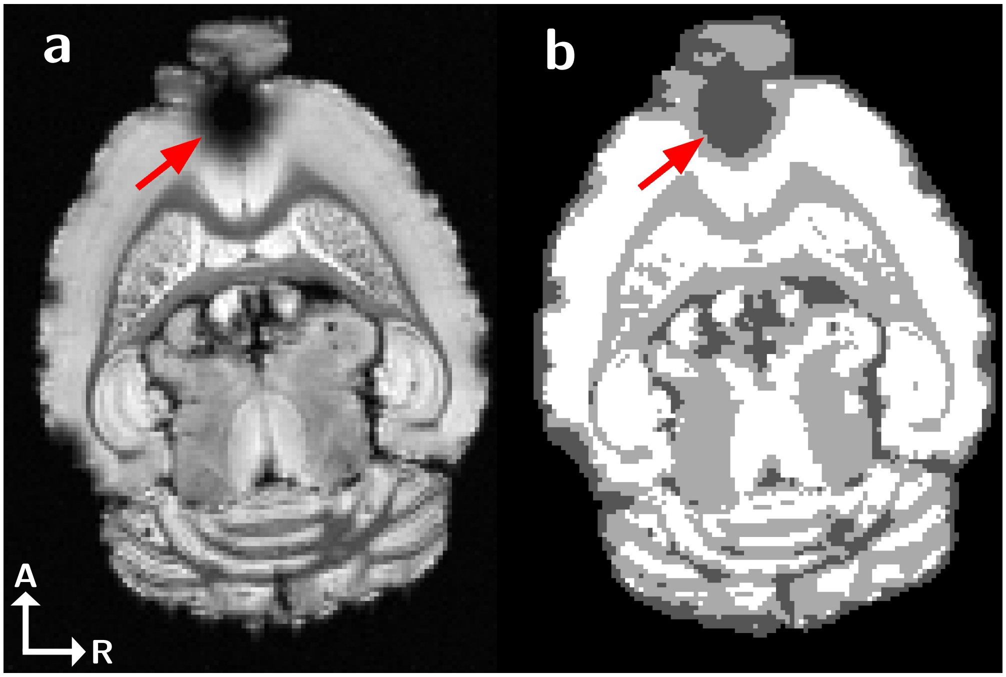


Figure S2: Demonstration of `Atropos` tissue segmentation results. Voxels in the CSF/susceptibility artifact class (dark gray, indicated by red arrow) were excluded from all analysis. A–R axis labels correspond to the Anterior and Right directions, respectively.

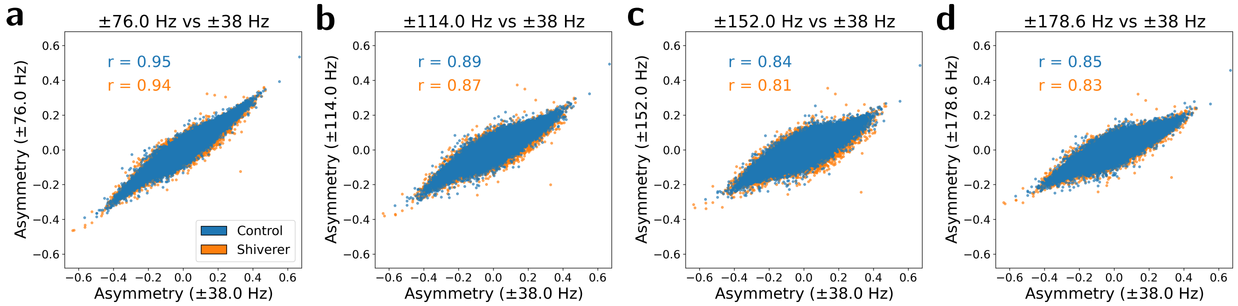


Figure S3: Scatterplots of data-derived asymmetry values calculated with a cutoff frequency of ±38 Hz vs. (a) ±76 Hz, (b) ±114 Hz, (c) ±152 Hz, and (d) ±178.6 Hz. r-values indicate Pearson's correlation coefficients, which remain above 0.8 for both control and shiverer datasets out to ±178.6 Hz.

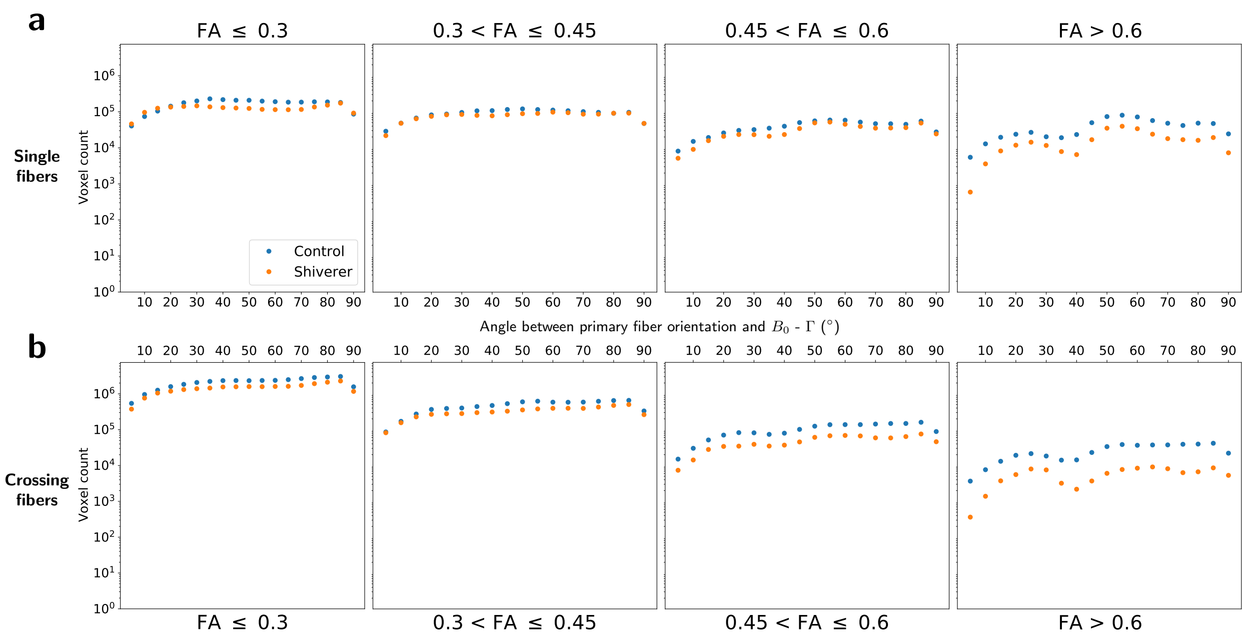


Figure S4: Voxel counts for control (blue) and shiverer (orange) data across FA and angular bins for voxels with (a) single and (b) crossing fiber populations.

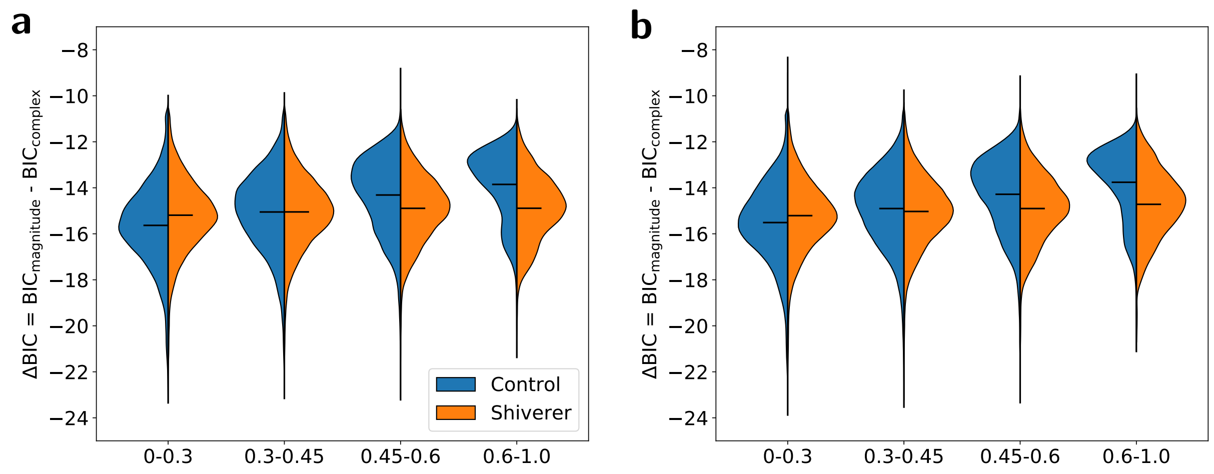


Figure S5: Distributions of absolute differences in BIC between the magnitude- and complex-fit models, separated by control/shiverer, FA bin, and (a) single and (b) crossing fiber voxels. The magnitude-fit model led to a consistently lower BIC than the complex-fit model.

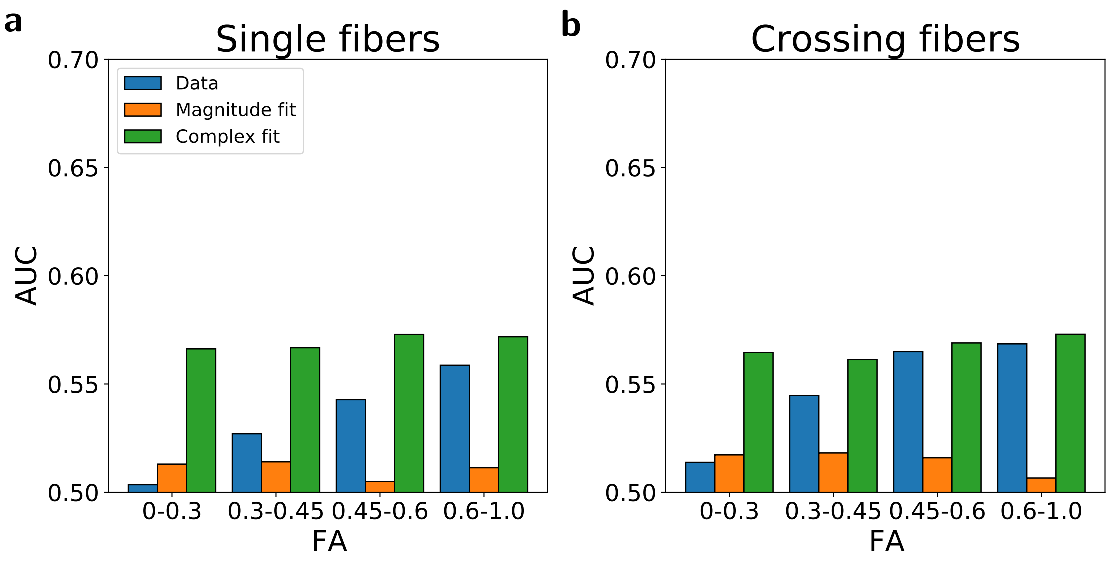


Figure S6: Values for the area under the ROC curve (AUC) using asymmetry as a one-variable classifier for control vs. shiverer data. FIDs were first truncated to 32 echoes prior to model-fitting and calculation of spectral asymmetry. Values represent AUCs for subsets of voxels in different FA bins containing either (a) single or (b) crossing fiber populations. With 32 echoes, neither the data nor either of the models is able to demonstrate meaningful sensitivity to myelin from the asymmetry, highlighting the need for high spectral resolution, or equivalently, extended FID sampling.

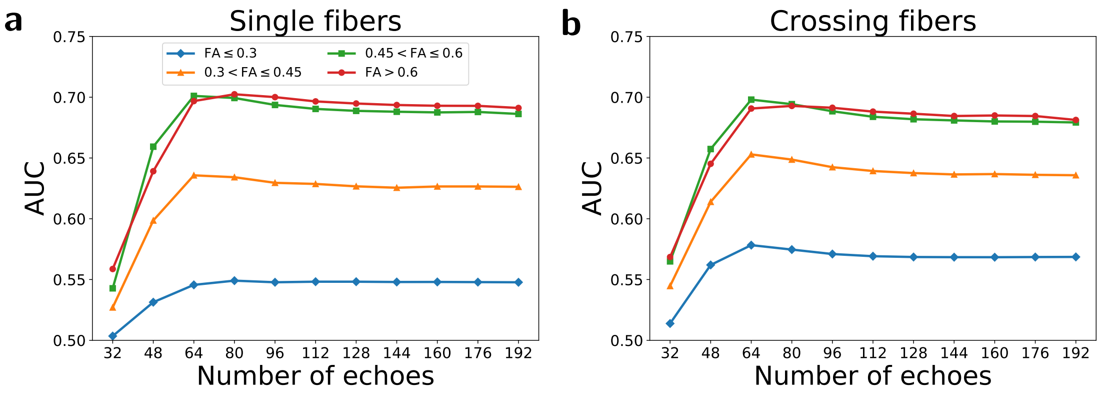


Figure S7: AUC values using data-derived asymmetry as a one-variable classifier for control vs. shiverer data as a function of the number of echoes in the FID. Subsampled-FIDs were created by truncating the full (192 echo) FIDs to the specified values prior to calculating asymmetry in the frequency domain. Values represent AUCs for subsets of voxels in different FA bins containing either (a) single or (b) crossing fiber populations. This demonstrates the importance of spectral resolution, or equivalently, extended FID sampling, and provides a roadmap for future benchmarking studies using EPSI spectral data.

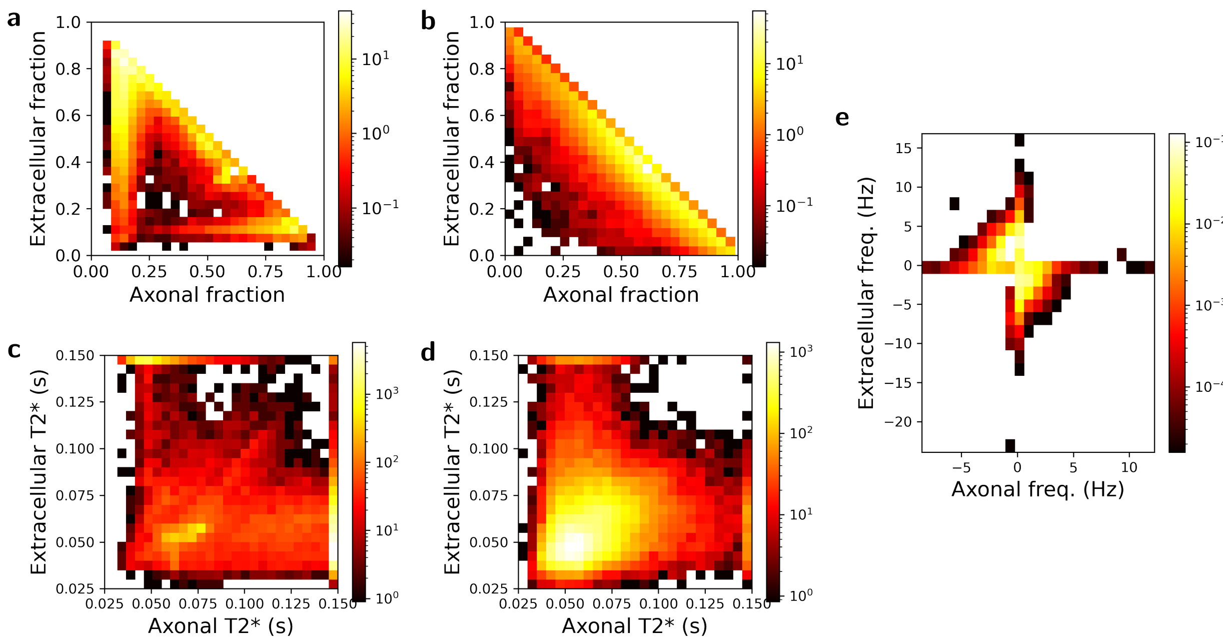


Figure S8: 2D distributions of model parameters for the "axonal" and "extracellular" components of both models for voxels with FA $> 0.6$. (a–b) Distributions of relative component amplitudes for the magnitude (a) and complex (b) models. (c–d) Distributions of T2* values for the magnitude (c) and amplitude (d) models. (e) Distributions of frequency shifts for the two components in the complex model. Note that for the magnitude model, there are a high number of voxels dominated strongly by the "extracellular" component over the axonal. Likewise, in the complex model, there are a high number of voxels in which the two compartment have similar T2* values and antisymmetric frequency shifts, consistent with lower spectral asymmetry values.
